# GeDex: A consensus Gene-disease Event Extraction System based on frequency patterns and supervised learning

**DOI:** 10.1101/839704

**Authors:** Larisa M. Soto, Roberto Olayo-Alarcón, David Alberto Velázquez-Ramírez, Adrián Munguía-Reyes, Yalbi Itzel Balderas-Martínez, Carlos-Francisco Méndez-Cruz, Julio Collado-Vides

## Abstract

**Motivation:** The genetic mechanisms involved in human diseases are fundamental in biomedical research. Several databases with curated associations between genes and diseases have emerged in the last decades. Although, due to the demanding and time consuming nature of manual curation of literature, they still lack large amounts of information. Current automatic approaches extract associations by considering each abstract or sentence independently. This approach could potentially lead to contradictions between individual cases. Therefore, there is a current need for automatic strategies that can provide a literature consensus of gene-disease associations, and are not prone to making contradictory predictions.

**Results:** Here, we present GeDex, an effective and freely available automatic approach to extract consensus gene-disease associations from biomedical literature based on a predictive model trained with four simple features. As far as we know, it is the only system that reports a single consensus prediction from multiple sentences supporting the same association. We tested our approach on the curated fraction of DisGeNet (f-score 0.77) and validated it on a manually curated dataset, obtaining a competitive performance when compared to pre-existing methods (f-score 0.74). In addition, we effectively recovered associations from an article collection of chronic pulmonary diseases, and discovered that a large proportion is not reported in current databases. Our results demonstrate that GeDex, despite its simplicity, is a competitive tool that can successfully assist the curation of existing databases.

**Availability:** GeDex is available at https://bitbucket.org/laigen/gedex/src/master/ and can be used as a docker image https://hub.docker.com/r/laigen/gedex

**Supplementary information:** Supplementary material are available at **bioRxiv** online.

## Introduction

Elucidating associations between genes and diseases is an important piece of information for genomic medicine. These associations help to understand the genetic mechanisms underlying diseases, contributing to the identification of targets and treatments. This knowledge serves as a basis for the development of gene therapies, an essential boost in genomic medicine.

For decades, several databases have manually curated and made available gene-disease associations from biomedical literature. Among these databases, one of the oldest is the Online Mendelian Inheritance in Man^®^ (OMIM^®^, https://omim.org/), which includes information about genes associated with Mendelian disorders. Another database is the Human Gene Mutation Database (HGMD^®^, http://www.hgmd.cf.ac.uk/ac/index.php), which reports associations between gene mutations and human inherited disease. Besides, the Comparative Toxicogenomics Database (ctd, http://ctdbase.org/) contains manually curated gene-disease associations, together with others like chemical-gene and chemical-disease relations.

As manual curation of gene-disease associations from the literature is time-consuming and demanding, several automatic approaches have been proposed. The most common and simple one is associating all the genes and diseases that are mentioned in the same sentence (i.e., co-occurrence). An improvement of this approach comes from methods like DISEASES (https://diseases.jensenlab.org) and DTMiner (Xu et al., 2016). The former exploits a weighted co-occurrence scoring scheme (Mørk et al., 2013; Pletscher-Frankild et al., 2015). The latter implements a binary Support Vector Machine (SVM) to determine if a gene-disease co-occurrence is a true association (https://gdr-web.rwebox.com/public_html/index.php). This SVM classifier employs the so-called local lexical features, which comprise words surrounding the gene and the disease. In addition, it incorporates global syntactic features such as unigrams, bigrams and trigrams of lemmas from the shortest path between a gene and a disease, and from the path between the least common ancestor of both entities and the root of the dependency tree (global syntactic features) (Xu et al., 2016).

The database DisGeNET (http://www.disgenet.org/) makes use of BeFree (Bravo et al., 2015), a system based on the combination of shallow features (lemmas, part-of-speech tags) with syntactic features (dependency tags) combining a Shallow Linguistic Kernel (Giuliano et al., 2006) with a Dependency Kernel (Kim et al., 2007). Recently, Deep Learning approaches have been applied to gene-disease association extraction, as in the case of ReNet (Wu et al., 2019), which obtains sentence representations from word embeddings using a convolutional neural network. Then, sentence representations are transformed into document representations using a recurrent neural network.

Despite these advances, extracting associations between genes and diseases from biomedical literature remains a challenging active research issue. Approaches extracting associations at sentence level propose unconfirmed, or hypothetical associations. Furthermore, these approaches could suggest contrary associations obtained from one sentence affirming the association and from another one rejecting it.

To face this challenge, some methods incorporate strategies to consider information beyond the sentence level. For example, DISEASES takes into account the frequency of the co-occurrence over a set of abstracts. DTMiner includes a weighting factor based on a citation network constructed from PubMed, which considers the relevance of the article where the association is reported. Also, DTMiner bears in mind if an author mentions the same association in several articles. More recent approaches take into account document-level associations, such as ReNet. These approaches aim to incorporate all the sentences where the same entities co-occur to make a single consensus prediction; instead of assigning a different class for each appearance of the association.

Another critical challenge for the extraction of associations is the scarcity of easy-to-use systems for curators, and the lack of helpful outputs to be utilized by them to feed databases. Altogether, this has led to a scarce representation of several genes and diseases in state-of-the-art databases, as shown by others (Bravo et al., 2015; Kim et al., 2017), further complicating their study by researchers of different fields. In this work, we tried to tackle this problem by developing a system that uses as input only a list of PubMed IDs, and returns associations in a curation-oriented format.

Here, we present Gene-disease Event Extraction System (GeDex), a freely available tool that extracts consensus gene-disease associations from biomedical literature with high performance. We followed the assumption that true associations are recurrently mentioned across the literature; a feature that could be exploited to distinguish true associations from false positive ones. This idea has been explored before using a scoring scheme that incorporates sentence and abstract-level co-occurrence to classify associations (Pletscher-Frankild et al., 2015). But as far as we know, GeDex is the first method that classifies gene-disease associations by taking into account all the available evidence supporting them.

Our method employs four simple features to classify each association: the number of mentions in the corpus (frequency) of the gene, of the disease, of the association itself, and a probability score that integrates those values. As we incorporate all the sentences that mention each association into these four features, we can make a single prediction that depicts the literature consensus. The individual distributions of these features do not show a clear separation of classes (true and false). Thus, using statistical methods, such as cut-off values, to separate true associations from false ones was not possible (Supplementary Figure S1). Therefore, we decided to apply supervised learning algorithms to find the underlying patterns that could separate between the two classes.

To test this, we used the number of mentions of genes, diseases, and their co-occurrence in a corpus of 11,721 abstracts, to train a random forest classifier (Breiman, 2001). The training dataset comprised 16,441 unique gene-disease co-occurrences extracted from the manually-curated fraction of DisGeNet. We achieved an f-score for the positive class of 0.76 in the testing stage. This score was robust despite variations in the training corpus composition, according to the results obtained in 100 independent training and testing runs (average f-score = 0.77, s.d. = 0.0072). We confirmed that a true pattern can be recognized from entity frequencies given that the model performance is consistently higher when compared to randomly generated frequencies. Moreover, in an evaluation using a manually curated small corpus with only 1145 unique associations, GeDex accomplished an f-score of 0.74. This fact confirms that our approach maintains its performance in smaller datasets.

GeDex was employed to extract associations of 433,475 articles related to chronic pulmonary diseases, aiming to assist the curation of a specialized database in the future. We observed that the vast majority of the predicted associations are not contained in current databases. This highlights the need to integrate automatic approaches into database curation, as the amount of published scientific articles increases faster than to cope with it manually.

## Methods

GeDex comprises five main stages (Figure 1), described in the following subsections. From a PubMed ID list, the corresponding abstracts were automatically downloaded using PubTator, which internally performs the Named entity recognition task to obtain annotated gene and disease mentions in the sentences of these abstracts (Annotation). Then, all gene-disease pairs are retrieved from each sentence, and entity mentions are normalized to identifiers (Association extraction). Afterwards, frequencies of genes, diseases, and their co-occurrence are calculated and scaled to be employed as features (Feature extraction). Next, a predictive model (classifier) assigns a class (true or false) to each association(Prediction). In the end, GeDex delivers these predictions along with their corresponding synonyms and supporting sentences (Enriched output).

**Fig. 1.**
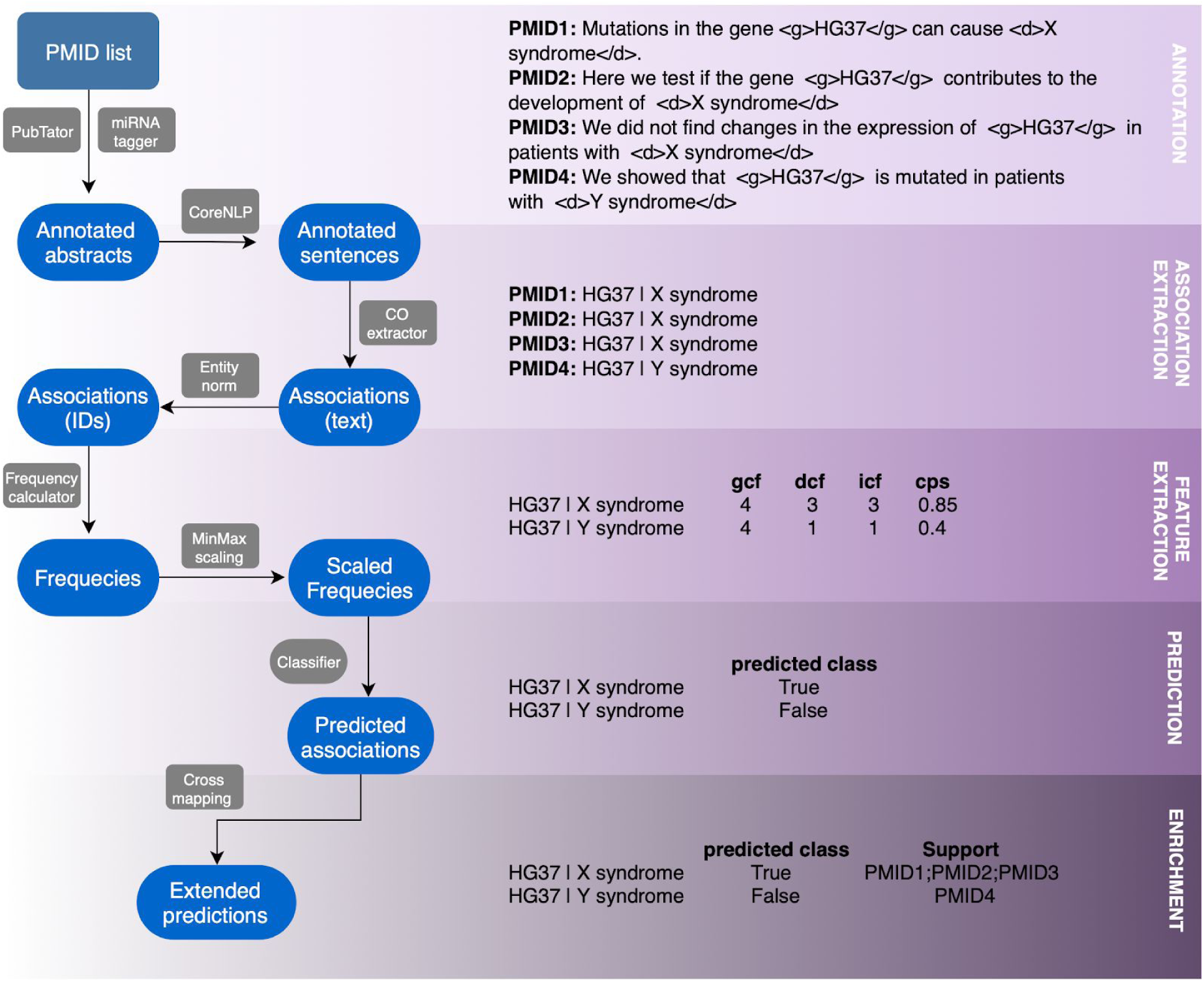
GeDex Pipeline. Our method comprises five main steps (green boxes) described in the Methods’ subsections. Blue ovals represent output files, and gray boxes are processing steps. The right panel depicts mock example of what the output of each stage would look like. We omitted the conversion of names to identifiers for simplicity and interpretability of the example. Traditional methods would return three different predictions for HG37 and X syndrome, one for each abstract that mentions it. As the example shows, the prediction for PMID1 and PMID2 would be that the association is true; but for PMID3 it would be false. From a curator’s perspective, this would be contradictory. As it is shown at the end of the example, our method solves this problem by returning a single prediction for the association.

### Data

To develop our method, we made use of the manually-curated fraction of DisGeNet v5.0 (Piñero et al., 2015) that includes manually-curated associations from ORPHANET (Pavan et al., 2017), PSYGENET (Gutiérrez-Sacristán et al., 2015), UNIPROT (Consortium et al., 2018) and CTD (Davis et al., 2018). This dataset included 11,721 abstracts with 16,441 non-redundant gene-disease associations. All the interactions were considered positives and therefore used as a gold standard. This means that any association not included in this dataset was considered negative. We divided this dataset in 80% for training-development and 20% for evaluation.

For the benchmarking procedure, we used a small corpus of pulmonary diseases manually curated by a research group in chronic pulmonary diseases from the National Institute of Respiratory Diseases of Mexico. This dataset comprised only 354 abstracts with 3361 text-based gene-disease associations. This was approximately 30 times smaller than the training dataset, resembling a more common situation for assisted curation projects. This dataset allowed us to observe if our approach performs well with small amounts of data.

The last dataset with 433,475 articles was utilized to extract gene-disease associations using GeDex. This dataset was delivered by the same research group to eventually curate and use the extracted associations for future research in pulmonary diseases. After extracting associations, we performed a database interrogation to discover those that have not been previously gathered in databases.

### Annotation

We used the PubTator RESTful APIs (Wei et al., 2013, 2012a,b) to obtain abstracts with annotated genes and diseases together with their respective NCBI and Medical Subject Headings (MeSH) identifiers. Additionally, we developed a set of regular expressions to annotate microRNAs (miRNA tagger).

Each abstract was split into individual sentences using the command line implementation of the Stanford CoreNLP version 3.9.1 (Manning et al., 2014). From these sentences, we selected only those containing at least one gene-disease co-occurrence for the remaining steps of the pipeline; conforming our corpus.

#### Association extraction

To extract all the existing associations between genes and diseases, we generated (CO extractor) all the possible gene-disease pairs using co-occurrence at the sentence level (text associations). Then, in order to facilitate a correct entity frequency calculation and a fairer comparison to reference databases, we normalized (Entity norm) gene and disease names to their respective NCBI and MeSH identifiers (Lipscomb, 2000) (ID associations), respectively.

During training, the DisGeNet disease identifiers were determined by utilizing the CTD disease vocabulary (MEDIC), as provided by the Dnorm 0.0.7 downloadable software (Leaman et al., 2013). Gene identifiers were kept as provided. Genes that shared the same identifier or had identical names were considered as synonyms. These synonyms were mapped to a reference DisGeNet identifier when possible, otherwise, the original identifier was conserved (Supplementary Figure S2). This procedure was necessary since DisGeNet assigns a unique identifier to homologous genes, almost always using the human identifier, while PubTator returns different identifiers for homologous genes. Entities that were annotated as gene or disease, but did not have an assigned identifier, were discarded and not considered during the entity frequency calculation.

To achieve a fair comparison to DisGeNet, we eliminated interactions from the database in which the gene or disease was not annotated in the abstract. This could occur when the association was curated from the body of the article and not directly from the abstract.

When analyzing a new corpus, GeDex considers genes with identical names, and/or identical identifiers, as synonyms and assigns a common internal identifier that is used for entity frequency calculation.

### Method development settings

A supervised learning approach can comprise different combinations of building blocks for each learning stage. It is a common practice to test all possible combinations of several elements guided by a rationale according to a particular task. To find the best-performing method, we tested three variations at the level of features, along with three methods to incorporate abstract-level features and two for corpus-level features; four feature scaling approaches and four different classifiers. This resulted in 128 different settings for the training stage (Supplementary Figure S3).

As we were exploring the reach of entity frequencies for classification of associations, we tested three different sets of features: only entity frequencies at the abstract-level (APS), only frequencies at the corpus-level (CPS), and using both simultaneously (BTH). We also tested three approaches to represent the abstract frequencies in the types aps and bth: all the values, their mean, and their median. In the BTH case, when all the abstract frequencies were used, we repeated the corpus frequencies; because this value is the same regardless of the abstract in which the co-occurrence appears. For the corpus frequencies, we tested two strategies: all values and consensus values. When all the values were included, the same corpus frequency was assigned to each PMID in which the co-occurrence appeared; we call this a triplet-based interaction (PMID-gene-disease). For the consensus values, the PMIDs associated to the co-occurrence were discarded, and the corpus frequency of the co-occurrence appears just once in the training data, as well as the pair gene-disease; we call this a tuple-based interaction (gene-disease). Using the corpus-level features of tuple-based interactions significantly outperformed the remaining settings (see, for example, dt-tup-cps and rf-tup-cps in Supplementary Figure S3).

To make GeDex robust and applicable to datasets of varying sizes, we tested three different scaling methods implemented in the scikit-learn machine learning tool (Pedregosa et al., 2011): normalization, robust scaling, and min-to-max scaling. We also tested the original values without applying any normalization or scaling method. Out of those, min-to-max scaling proved to be the best when combined with the other top-performing components (features and classifier) (Supplementary Figure S3).

To find the best logic to disentangle the complex patterns of entity frequencies (Supplementary Figure S1), we also tested four of the classifiers implemented in scikit-learn: multinomial Bayes, random forest, decision tree, and logistic regressor. Out of those, the random forest classifier was among the top performers (see Supplementary Figure S3 and section). Therefore, in GeDex we implemented a random forest classifier trained on the corpus frequencies scaled using the method min-to-max.

### Feature extraction

#### Frequency calculation

For each entity (gene, disease, and co-occurrence), we compute the number of mentions per abstract and then add them to get the overall frequency in the corpus, as follows:

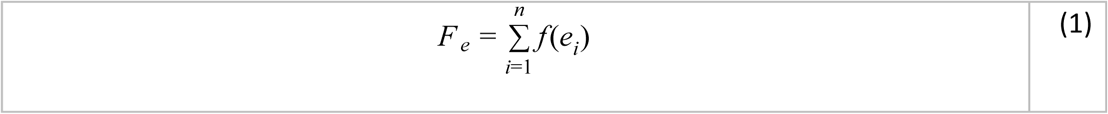

where *f* (*e*_*i*_) is the frequency of the entity *e*_*i*_ in the abstract *i*; *F*_*e*_ is the corpus frequency of the entity *e*; and *n* is the total number of abstracts.

As a fourth feature, we designed a simple probability score to reflect the relevance of the co-occurrence given the corpus frequency of the gene and the disease comprising it. The score is calculated as follows:

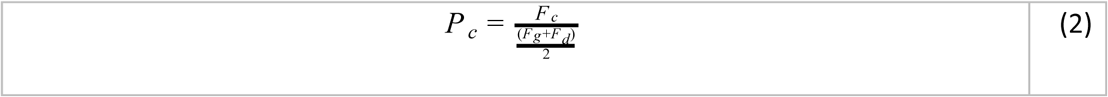

where *P*_*c*_ is the probability score, *F*_*c*_ is the corpus frequency of the co-occurrence, *F*_*g*_ is the corpus frequency of the gene, and *F*_*d*_ is the corpus frequency of the disease. For example, if a gene and a disease appear 100 times in the corpus, and co-occur 100 times, this score gives 1.

#### Min-to-Max Scaling

This method constrains the values to a specific fixed range. As previously stated, we use this method to deal with variations in the corpus size. The scaled values are calculated as follows:

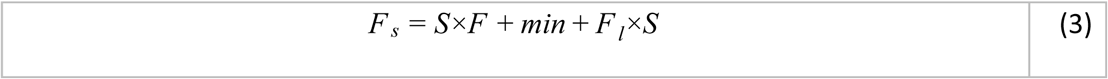

where

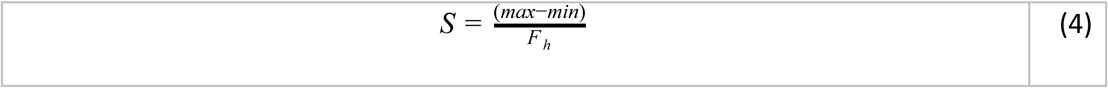

where *F* _*s*_ is the vector of scaled values per feature; *F* is the vector of original values per feature; *min* and *max* are the desired minimum and maximum of the scaled values, respectively (in this case 0 and 1); *F*_*l*_ and *F*_*h*_ are the minimum and maximum of the original feature values, respectively.

### Prediction

#### Supervised learning approach

Here, we solved our problem as a binary classification task, where the two possible classes for any given gene-disease association are positive and negative. Our predictive model was a random forest classifier trained using the four features described in the previous section. A random forest classifier is a model that integrates several decision trees fitted to subsamples of the original training dataset and calculates the average of their individual probabilities for each prediction to make a final decision (Breiman, 2001). We used the default parameters of the scikit-learn implementation, which uses ten random trees without constraining their maximum depth and uses bootstrap to sample the observations used to build each tree (Pedregosa et al., 2011). Also, we used the *balanced* mode, which automatically weights each class based on its frequency in the training data, as there was a slight class imbalance in our training dataset (Supplementary Figure S4). As we employed 100 validation folds to score the performance of our predictive model, we confirmed that each fold had approximately a 25% of unique associations in comparison with all remaining folds (Supplementary Figure S4).

### Enriched output

GeDex provides a helpful output, designed to facilitate manual validation efforts. To this end, each gene-disease association is reported with its predicted class, along with all evidence sentences found in the complete article collection and the PMIDs of the corresponding articles. Also, GeDex provides all the names found for a given gene and disease, as well as synonymous gene identifiers.

### Evaluation

To asses GeDex’s performance, we selected five widely used metrics: precision, recall, f-score, area under the receiver operating characteristic (AUROC), and area under the precision-recall curves (AUPRC). As there is no reference for true negative gene-disease associations, we evaluated the performance of the positive class. This approach was maintained throughout the testing, validation, and benchmarking steps.

Precision refers to the ratio between the total number of associations retrieved that are true positives and the overall number of instances retrieved as positive. Recall is the total number of true positive instances retrieved, divided by the total number of instances in the gold standard (Ting, 2010). The harmonic mean of these two metrics gives the f-score (Sammut and Webb, 2010b). The ROC curve is a visual analysis of the relationship between the sensitivity (or true positive rate) and the specificity (or false positive rate) (Sammut and Webb, 2010c), while varying the threshold value for positive classifications from its minimum to its maximum (decision threshold) (Sammut and Webb, 2010a). An AUROC greater than 0.5 indicates that a classifier performs better than randomly assigning classes. On the other hand, the PRC displays the trade-off between precision and recall at different decision thresholds (Ting, 2010).

#### Training-testing runs

We compared our results to a simple co-occurrence approach as a baseline method. To discard biases due to dataset composition, we performed 100 training-testing independent runs (development folds) by randomly sampling 60% of the interactions in the development dataset (Supplementary Figure S4a) and using the evaluation fraction (See Data section of Methods) as testing dataset. Additionally, we generated synthetic frequencies using a uniform probability model constrained by the minimum and maximum values of the observed frequencies. We also performed 100 development folds on this dataset. Both the observed folds and the random folds had an average of 25% unique associations when compared to the rest of the folds accordingly (Supplementary Figure S4b).

#### Validation and benchmarking

To compare GeDex against a human reader, we assessed our algorithm’s performance using a pre-existing collection of abstracts that had been manually curated by a research group interested in chronic pulmonary diseases. During curation, sentences were classified as negative instances when a gene-disease relationship is denied or is not explicitly mentioned. If a relationship is explicitly mentioned, then the sentence is classified as a positive instance. While this strategy is convenient for evaluating performance on specific sentences, it creates difficulties when considering consensus classifications, since a given gene-disease pair could be classified as positive and negative depending on the explicitness of the sentence.

Sentences with gene-disease co-occurrences, were classified as positive when an association between these entities existed, and as negative when the relationship is denied, or there is no mention of an association. This strategy generated examples where the same gene-disease pair had a positive and a negative classification during manual curation.

Therefore, to evaluate GeDex’s ability to predict associations, we created three types of consensus: a Positive Consensus, where gene-disease pairs with at least one positive classification were left as positive in the consensus; a Majority Consensus, where the class was decided by which classification was more common for the gene-disease pair; and a No Ambiguity Consensus by removing all gene-disease pairs that had more than one class.

For benchmarking, we selected previously described text mining systems that were freely available. DTMiner (we leveraged the implementation in ReNet’s benchmarking), BeFree (implemented at https://bitbucket.org/ibi_group/befree/src/master/), and ReNet(from https://bitbucket.org/alexwuhkucs/gda-extraction/src/master/) were applied and evaluated on the same validation corpus described previously. In contrast to GeDex, these systems were not designed to infer corpus consensus classifications. Instead, they classify on a case-to-case basis, analyzing each sentence or abstract separately. This leads to a similar situation described during manual curation, where gene-disease pairs have more than one class. To resolve this, we generated consensus predictions in the same way as described previously and compared the consensus predictions to their corresponding reference; Positive Consensus predictions, for example, are evaluated against a Positive Consensus reference.

One of the many challenges of comparing different tools was reconciling differences in recognized entities. This was particularly challenging when using BeFree, since this tool employs its own Named Entity Recognizer, while the other tools employ PubTator. To mitigate this situation, the evaluation of predictions was limited to entities that were mentioned in the manually curated reference. This, however, meant that many of BeFree’s predictions were not considered.

To evaluate class prediction bias, we sampled 50 subsets with a balanced number of positive and negative associations from the manually curated corpus. The GeDex pipeline was applied to these subsets, and predictions were generated for each sample.

### Database interrogation

We used the Phenotype-Genotype Integrator of NCBI (PhenGenI) (Ramos et al., 2014) and all the associations reported on DisGeNet v5.0. The former incorporates GWAS results with several databases, including Gene (Maglott et al., 2010), dbGap (Tryka et al., 2013), OMIM, and dbSNP (Sherry et al., 2001). DisGeNet bears a compendium of gene-disease associations from UNIPROT, CGI (Tamborero et al., 2018), ClinGen (Rehm et al., 2015), Genomics England (Caulfield et al., 2017), CTD (human subset), PsyGeNET, and Orphanet. To ease the comparison, gene names were normalized using NCBI gene identifiers, and disease names were normalized with the Medical Subject Headings (MeSH) identifiers.

For cases in which several gene IDs in our predictions were considered as synonyms (through gene names), all the combinations of synonym IDs were paired with the corresponding disease and searched in the reference databases. When the interactions were found in several databases, all of them were reported.

#### Implementation

GeDex is an automatic system that incorporates several available tools with home-developed programs. The whole system requires Java, Python v. 3.6, and R v. 3.5.0. All the scripts are organized within a shell script to ease its use. To run GeDex, only a list of PubMed IDs is required; see the documentation available at https://bitbucket.org/laigen/gedex/src/master/. GeDex can be cloned from the Bitbucket repository but is also available as a Docker image to avoid installing tools and libraries https://hub.docker.com/r/laigen/gedex.

## Results and discussion

### The supervised learning approach learns the literature consensus of associations using frequency patterns

GeDex aims to classify interactions based on the consensus of the sentences in the corpus, providing highly robust and coherent results. It employs the frequencies of genes, diseases, and their co-occurrence in the training corpus to classify each association. By doing this, we are taking into account the amount of evidence in the literature to support each prediction. Evidently, there is a dependency on the triage of the article collection to be curated, but this issue escapes the scope of our work.

Here, we show that frequency patterns can be captured by a supervised learning approach, and present a clear improvement compared to the simple co-occurrence approach. The random forest classifier consistently outperformed the co-occurrence approach (average f-score 0.57) on the testing step, with an average f-score of 0.77 on 100 development folds (Figure 2a). We achieved an average AUPRC of 0.82 and AUROC of 0.85 (Figure 2b), indicating that our method’s performance is maintained on both classes (Figure 2c,d), despite the imbalance of the training data (Supplementary Figure S4).

**Fig. 2.**
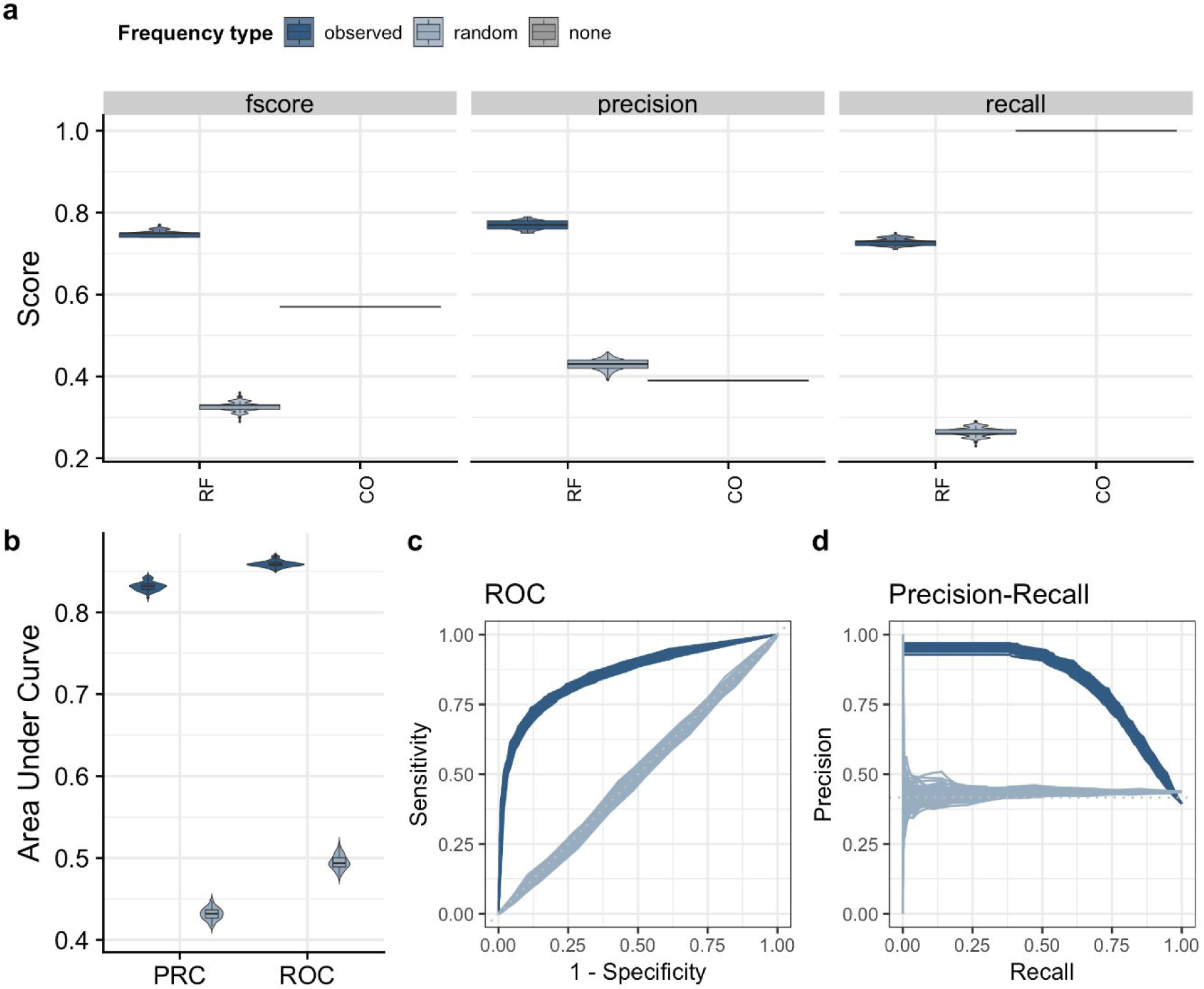
GeDex performance over 100 development folds. a) Distributions of precision, recall, and f-score for the co-occurrence approach (CO), the random forest classifier (RF) trained using observed and random frequencies. Mean precision = 0.78: mean recall = 0.76: mean f-score = 0.77. b) Distributions of the areas under the PRC and ROC curves for the RF classifier in both training settings (observed and random). c) ROC curves of the 100 development folds for the RF classifier. d) PRC curves of the 100 development folds for the RF classifier.

### Frequency patterns are independent of corpus composition

To test whether the corpus composition affected the frequency patterns, we generated 100 development folds (See Evaluation section in Methods) and measured the performance on each one (see Methods). The sampling procedure created folds with 25% of unique interactions on average (Supplementary Figure S4). We found very similar performance with all of them, as shown by the low variability on the performance metrics (f-score sd = 0.0057; Precision sd = 0.0077; Recall sd = 0.0073) (Figure 2a). The distribution of areas under the ROC and PRC curves for our classifier show very constrained ranges, indicating few changes in performance (Figure 2b). Therefore, on this dataset, the corpus composition did not affect GeDex’s performance.

### Entity frequencies show a true pattern and are not random

To show that our approach rescues a recognizable pattern in entity frequencies that distinguishes true gene-disease associations from false associations, we tested the performance of our model with 100 folds of randomly generated frequencies. This was achieved by sampling a uniform distribution limited by the highest and lowest of the observed frequency type (See Methods).

As shown in Figure 2a and b, the performance of the model is consistently and noticeably worse when trained and applied on random frequencies, compared to its performance using observed frequencies across all metrics. This shows that the observed frequencies comprise a representative pattern that is recognized by our classifier and serves to distinguish true positive associations from those that are false.

### GeDex performs similar to other methods in a manually-curated corpus

As stated, we leveraged a preexisting collection of abstracts manually curated by a research group. These manually classified interactions were aggregated into a consensus using three different strategies described in Methods. GeDex’s performance and benchmarking against BeFree, ReNet, and DTMiner can be observed in Figure 3. Here we show results only when evaluating predictions against the Positive Consensus. Results obtained with other Consensus are shown in Supplementary Figure S5.

**Fig. 3.**
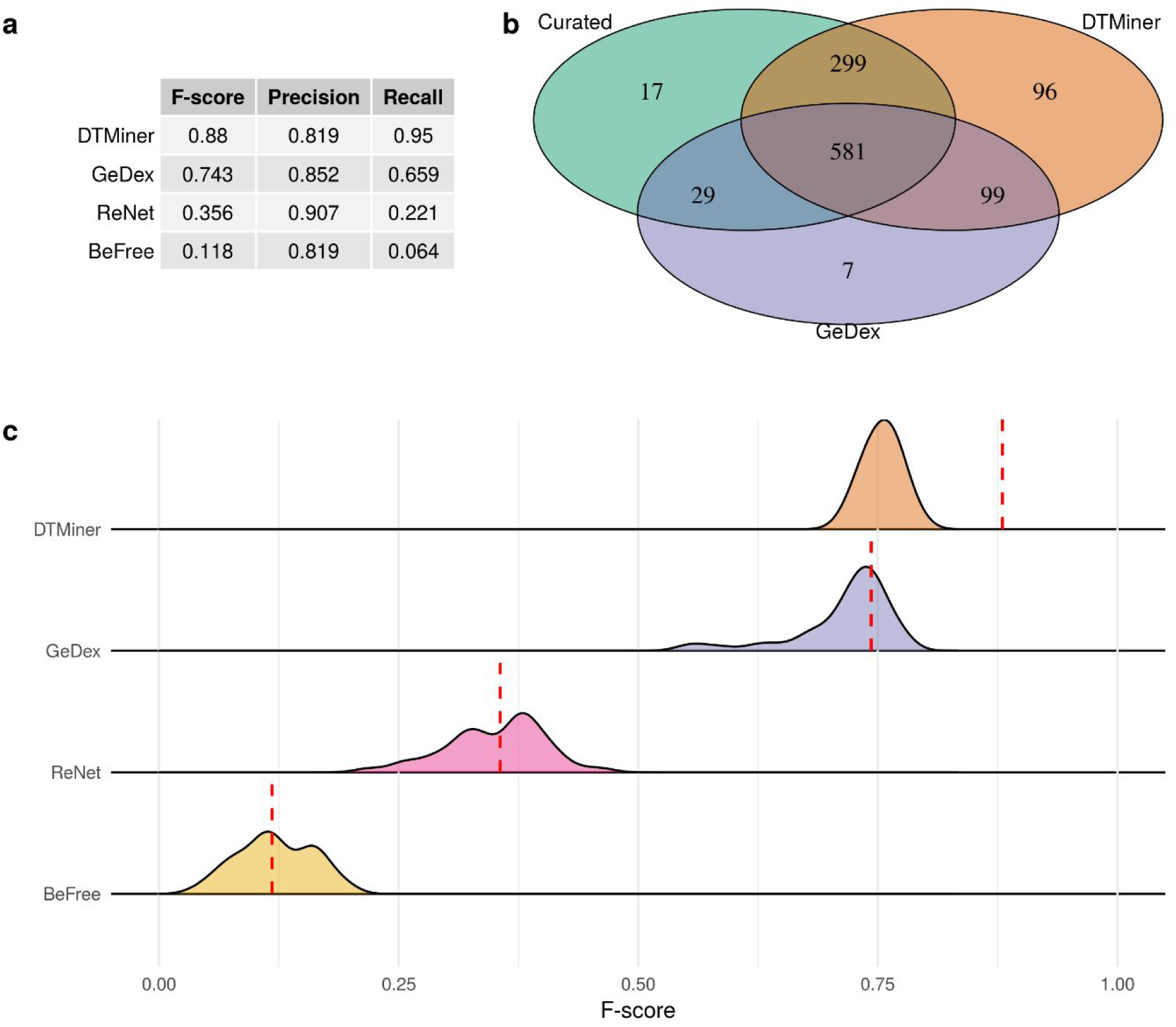
GeDex validation and benchmarking. a) Table of F-score, Precision, and Recall obtained by applying each tool to the manually-curated corpus. b) Venn diagram of positively classified associations between GeDex, DTMiner, and the human curator c) Distribution of F-scores obtained by applying each tool to class-balanced samples of the complete corpus (shaded area). The F-score obtained with the complete corpus is marked by a dashed red line.

We observed that GeDex’s performance does not vary significantly from the metrics obtained during testing. Remember that this manually-curated corpus is approximately 30 times smaller than the training dataset and, contrary to it, this contains a higher number of instances of the positive class than the negative one. This demonstrates that GeDex has a consistent performance in collections of different size and class distribution, supporting its application in different real curation scenarios.

When compared to other established text mining tools, GeDex performs competitively, showing improvement over BeFree and ReNet. It does, however, appear to have lower performance than DTMiner (Figure 3a). Even so, as can be seen in Figure 3b, DTMiner has a much higher number of false-positive classifications when compared to GeDex, resulting in our algorithm having a higher precision (Figure 3a). This led us to believe that DTMiner’s good performance was due to a heavy bias toward positive classifications. This bias would greatly improve the evaluated metrics since the curated corpus has a high number of positive gene-disease associations (Supplementary Figure S6).

When we evaluated against randomly selected associations, in a way that we obtained class balanced subsets of the complete corpus, it can be appreciated that DTMiner’s performance becomes comparable to that of GeDex (Figure 3c). Furthermore, DTMiner’s F-score distribution on these balanced subsets is distinctly inferior to its performance on the complete corpus (dotted red line in Figure 3c). This contrasted with GeDex, where performance on sampled subsets was not easily distinguishable from its F-score on the complete corpus (dotted red line), that is, the distribution of F-scores contains the score obtained when evaluated on the complete corpus. This result signals that GeDex is robust to class imbalance and delivers consistent results.

### GeDex retrieves associations related to pulmonary diseases not reported in databases

As a case study, we analyzed 433,475 abstracts extracted using the keywords Chronic Obstructive Lung Disease, Chronic Obstructive Pulmonary Disease, Combined Pulmonary Fibrosis and Emphysema, Idiopathic Pulmonary Fibrosis, Lung Cancer, and Pulmonary Fibrosis. We were able to extract sentences with gene-disease co-occurrences from 108,667 of them. We retrieved 792,597 text-based gene-disease associations, and we were able to normalize 786,886 of them. From an assisted curation perspective, this is a useful filter, as it significantly reduces the literature to be reviewed in downstream analysis.

To assess GeDex’s knowledge-discovery index, we searched the gene-diseases associations predicted as positive in several state-of-the-art databases (See section Database interrogation in Methods). We found that 70% of them were not reported in any of the searched databases (Figure 4a), further confirming that there is a vast amount of information present in the published literature that has not yet been reported in any database. As of the remaining 30% of our predictions, 20% were also reported by DisGeNet’s automatic method BeFree in the version v5.0 of the database, and the other 10% was found in the remaining databases (Figure 4a). The associations not reported in searched databases are available in the GeDex’s bitbucket repository.

**Fig. 4.**
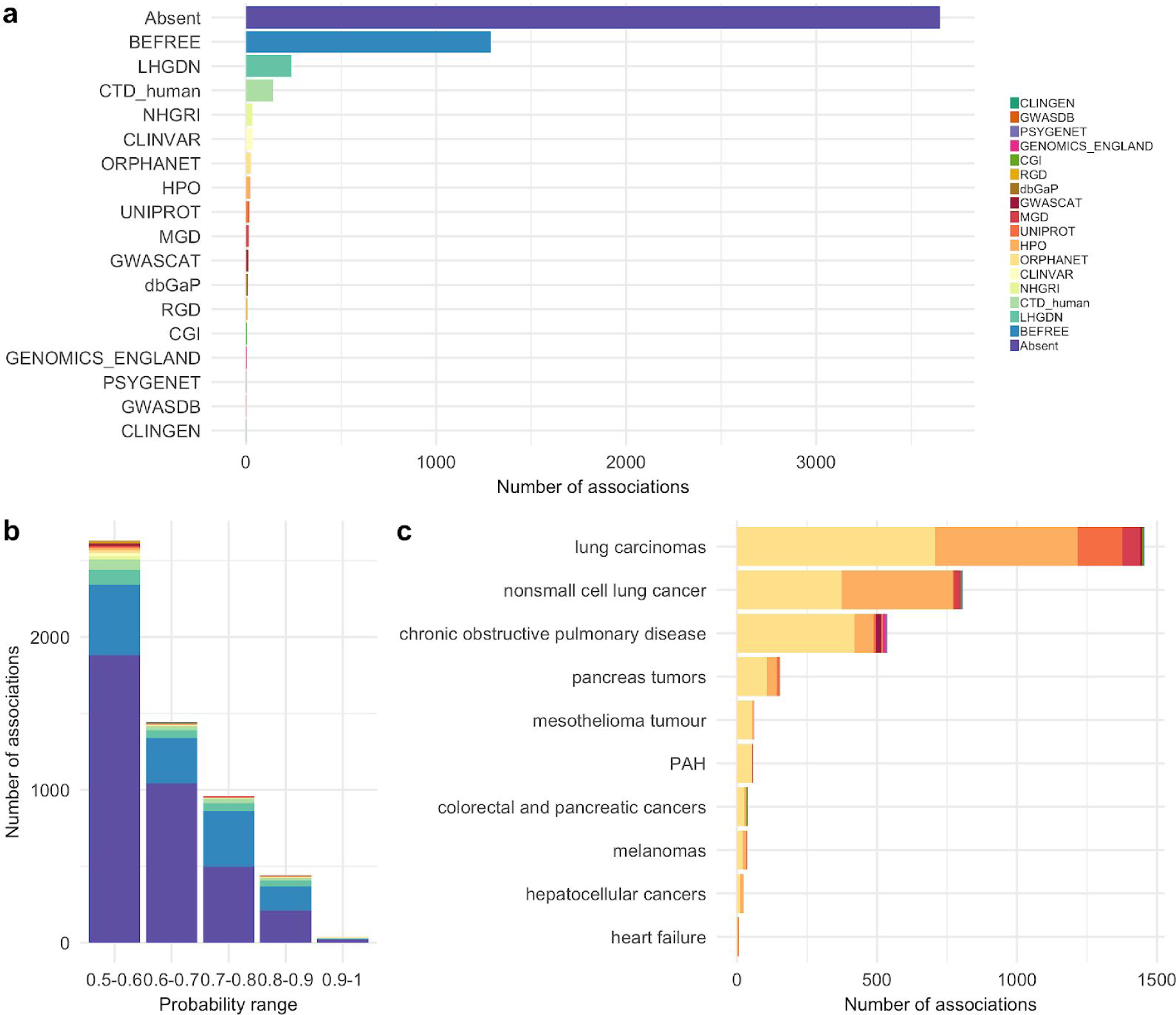
Database representations of gene-disease associations retrieved in our case study on pulmonary diseases. a) Source summary of gene-disease associations; the Absent source represents that the association was not reported in any of the databases searched. b) The proportion of sources grouped by the probability of positive class. c) Sources of gene-disease association of the ten most frequent diseases in our case study dataset.

Additionally, we analyzed the distribution of gene-disease association sources by their positive class probabilities (Figure 4b). We found that the highest number of positive interactions were within the range of 0.5 and 0.6 of probability. Given that this is the lowest probability range for positive interaction, the validity of the predictions in this range could be questioned. Although, the majority of predicted associations found in current databases were also within this range. We interpret this as a factor reinforcing the credibility of the absent associations that also lie within this low positive class probability region.

To further assess the validity of our predictions in this blind setting, we looked at the sources of associations comprised of the ten more frequent diseases in our data set (Figure 4c). We found that all of them were reported in at least one database. This also highlights the potential of GeDEx, as it was able to retrieve these expected associations, given the high amount of literature surrounding these diseases.

As a final control, we manually inspected the 60 gene-disease associations with the highest positive class probability and determined both their true class (based on the sentence) and the accuracy of their assigned sources. We noticed that genes and diseases often are not associated with all their synonyms in reference databases. To make a fair comparison, the manual classification was restricted to the identifier reported in the prediction (instead of the name). The most frequent errors were related to the annotation process; since 18 sentences had words wrongly tagged as genes or diseases by the NER system of PubTator.

These cases were mainly due to abbreviations or words that were similar to real names of genes and diseases. After discarding the annotation errors, we accurately classified 36 of 42 associations, achieving a precision of 0.85. Although the NER process is performed by an external tool, we are aware of the limitations it imposes for our method and any other text mining tools.

## Conclusion

Here we presented GeDex, a consensus Gene-disease Event Extraction System based on frequency patterns and supervised learning. GeDex obtained a competitive performance extracting associations from a manually-curated test corpus and against three available state-of-the-art gene-disease association extraction systems. In addition, we employed GeDex to extract association from an extensive article collection given by a research group in chronic pulmonary diseases from the National Institute of Respiratory Diseases of Mexico. Considering that many of these associations were not present in databases, and our competitive results, we see GeDex as a useful tool for gene-disease association extraction. With new studies being published at a brisk pace, it becomes difficult to maintain a completely up-to-date reference. Therefore, it will be interesting to note the fraction of gene-disease associations predicted by GeDex that will become present in future releases of the databases interrogated in our study, providing further validation for our tool. Furthermore, we recognise that newly associated genes have the potential to provide novel insights into the molecular mechanisms behind the diseases in which they are implicated. This could be addressed by analyzing the biological functions and biochemical pathways described for these gene groups.

Following this idea, it is important that automatic association extraction systems, such as GeDex, are easy to access and use. In the future, we seek to implement an online interface that can be employed for quick analysis and curation assistance. This would be very useful and valuable for scientists of diverse backgrounds and skill sets.

Finally, while we show that entity frequencies can be used to extract a consensus of gene-disease associations from literature, it will be interesting to explore the usefulness of this approach to analyse other types of biological associations. This will require, in part, a better understanding of the patterns learned by our algorithm. These patterns could, potentially, be employed to extract other types of associations.

## Acknowledgements

We acknowledge Alfredo José Hernández Álvarez and César Bonavides Martínez for computational support. Also, this work received support from Luis Aguilar of the Laboratorio Nacional de Visualización Científica Avanzada.

