## Supplemental File 1 for "GeDex: A consensus Gene-disease Event Extraction System based on frequency patterns and supervised learning" (from another version of this preprint: version 1, 2019-11-12)

### *GeDex Supplementary document*

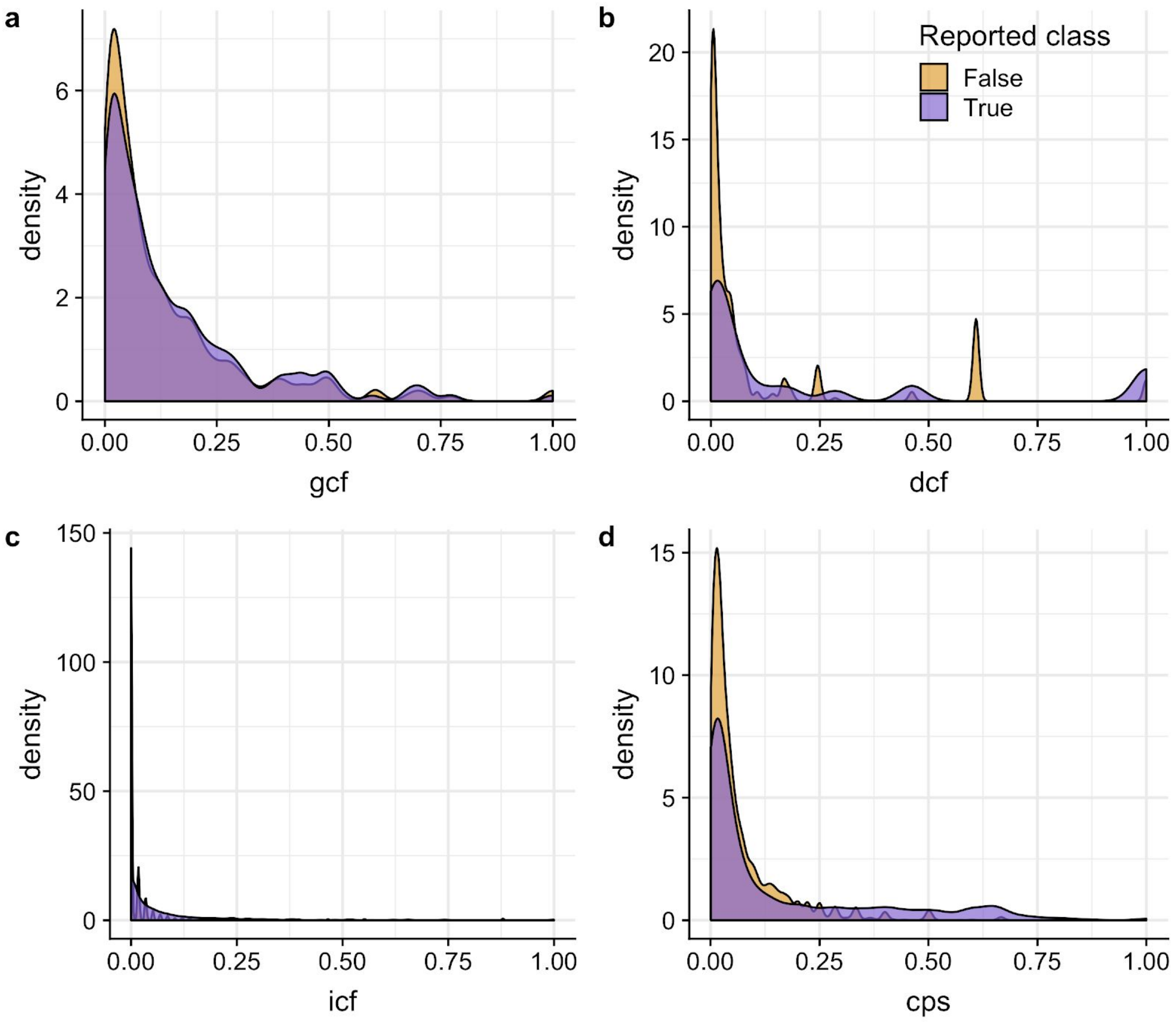


**Figure S1.** Distributions of min-to-max scaled values of the three observed features of the training data set. **a)** Distribution of gene corpus frequencies **b)** disease corpus frequencies **c)** frequency of their co-occurrence. **d)** Distribution of probability scores. gcf = gene corpus frequency; dcf = disease corpus frequency; icf = co-occurrence corpus frequency; cps = corpus probability score. The distributions are colored according to their true class (reported in DisGeNet), as shown in the upper panel legend.

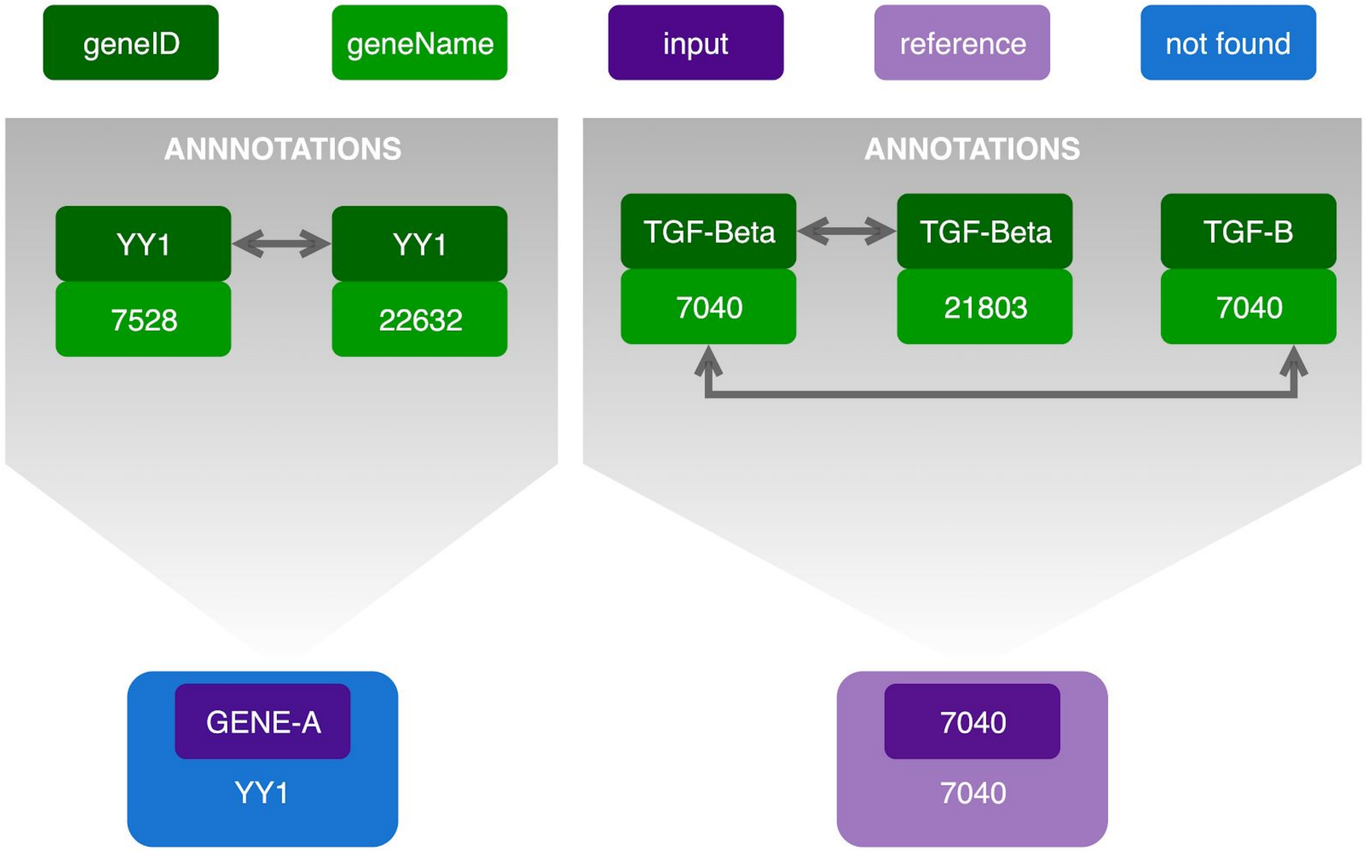


**Figure S2.** Entity normalization procedure. Here we show an example of how GeDex creates gene synonym groups. There are two ways in which synonyms are determined: one is by shared identifiers, as is shown for **TGF-Beta** and **TGF-B,** both having **7040** as their NCBI identifier. The other is by identical names; this can be seen for **TGF-Beta** entities, despite being homologous genes and having different identifiers, we consider them synonyms. Therefore both **TGF-Beta** entities and **TGF-B** are considered as the same gene with **21803** and **7040** as possible identifiers. When determining a common identifier during training, we searched for an identifier in the DisGeNet reference. In this example, all references to **TGF-Beta** in DisGeNet have the **7040** identifier. Therefore, during entity frequency calculations and evaluation, genes in the synonym group will have **7040** as their identifier. When analyzing a new corpus for predictions, or an identifier cannot be found in DisGeNet, an internally generated identifier is assigned. An example of this can be seen for **YY1**; since this gene is not present in DisGeNet, genes in this group are assigned an identifier based on alphabetic characters, so as to not be confused with NCBI numerical identifiers. This assignment also occurs when GeDex generated predictions for a new corpus.

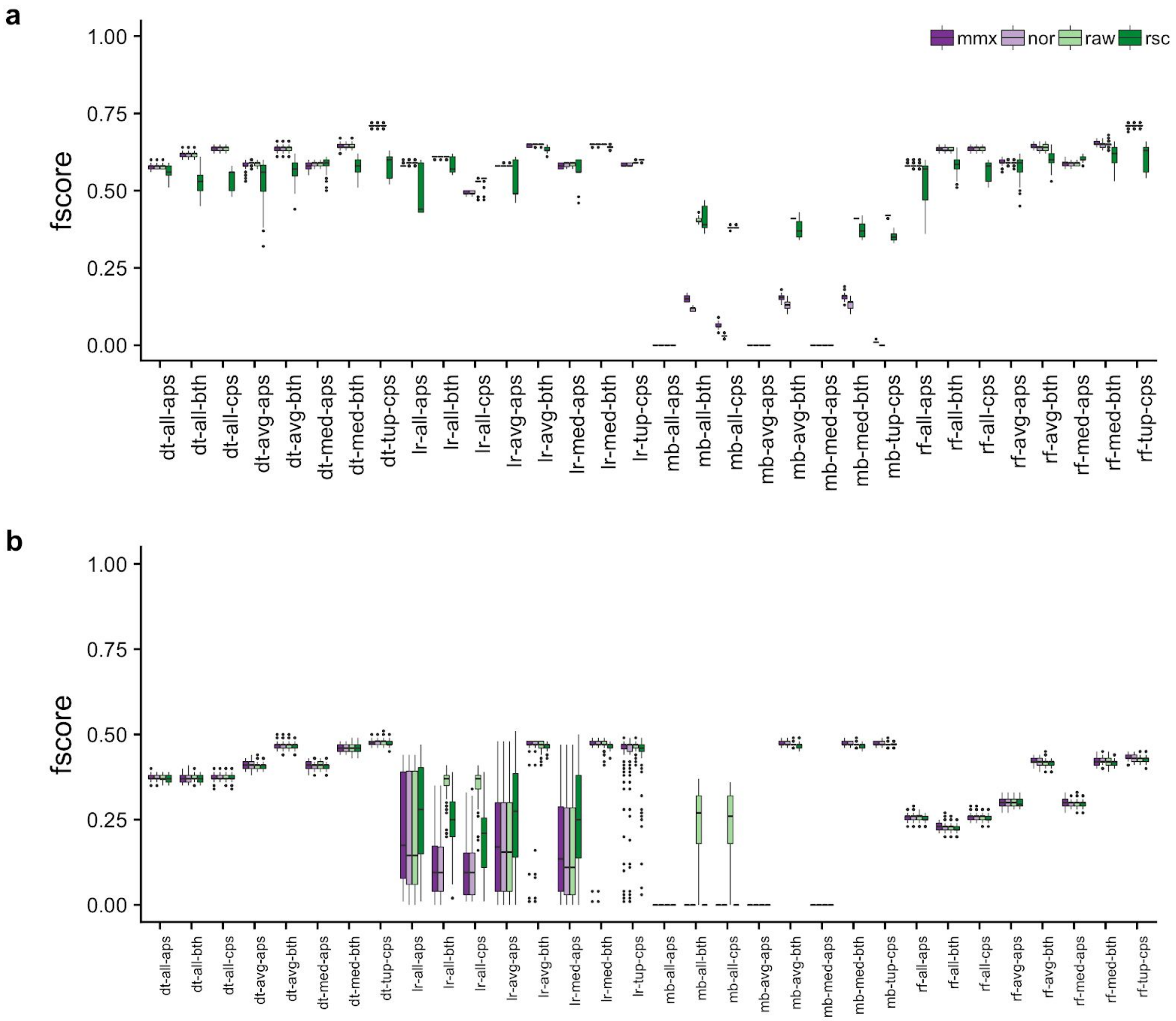


**Figure S3.** Performance of all combinations tested for method's development stage. The boxplots represent the distributions of f-scores over 100 training-testing runs using each set of elements. **a)** Performance of the classifier trained using observed frequencies. **b)** Performance of the classifier trained using random frequencies. The different colors represent the scaling method in both panels, as shown in the legend of the upper panel. Classifiers: lr = logistic regressor, dt = decision tree, mb =multinomial bayes, rf = random forest; Scaling: mmx = min-to-max, nor = normalization, rsc = robust scaler, raw = original values without scaling; Level of features: aps = abstract frequencies, cps = corpus frequencies, bth = aps and cps; Method to incorporate features: med = median integration; avg = average integration; all = all values included.

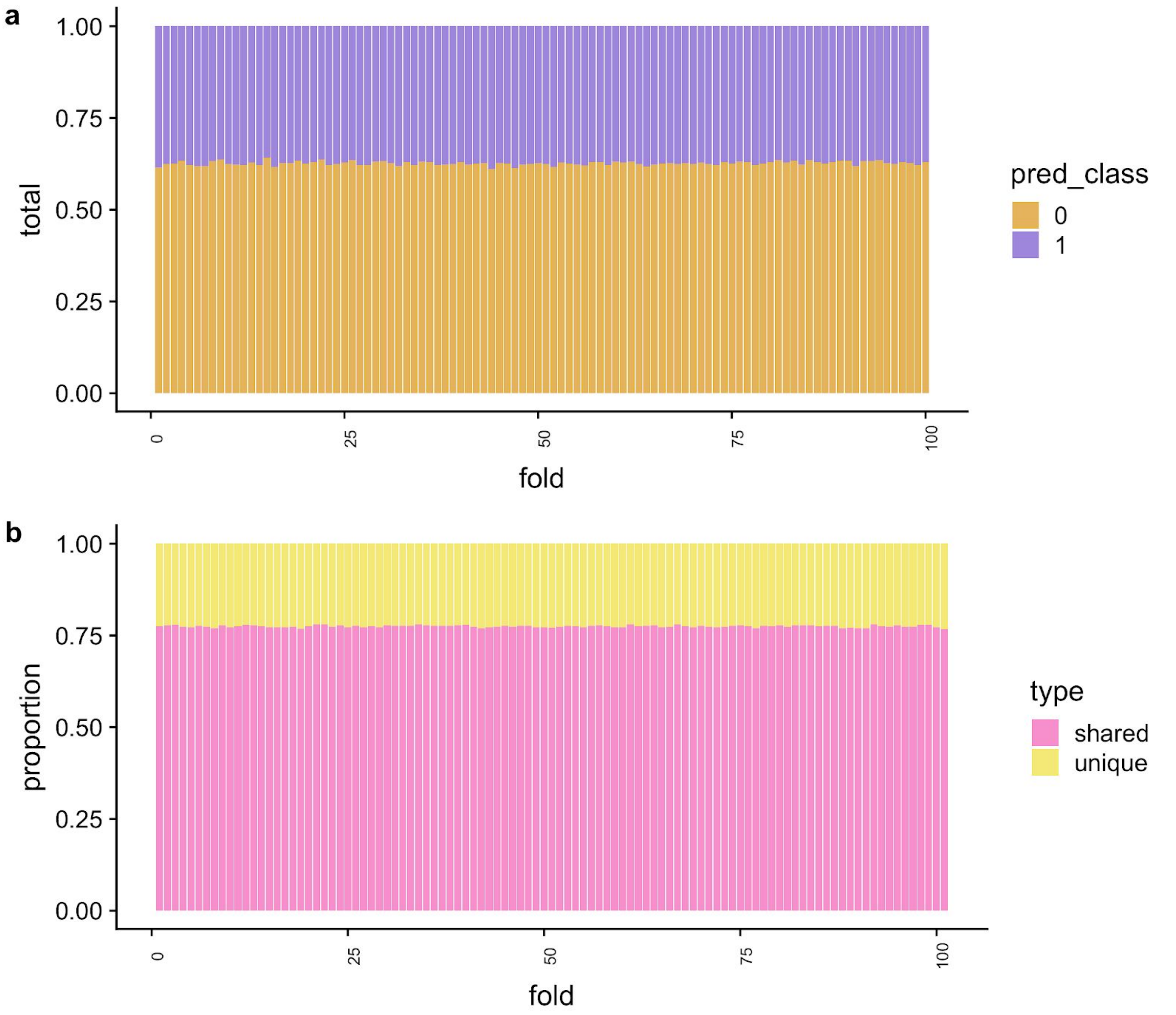


**Figure S4.** Class distribution and composition of training data sets of the 100 development folds. **a)** class proportions (y axis) of each fold (x axis) as shown in the legend (0 = negative clas; 1 = positive class). This shows imbalance data. **b)** Average proportion of associations shared with all other folds and unique to each fold, as shown in the legend. This shows that folds contain an approximately 25% of unique associations to train our classifier.

**Majority Consensus**

| | F-score | Precision | Recall |
|---|---|---|---|
| DTMiner | 0.819 | 0.743 | 0.913 |
| GeDex | 0.682 | 0.752 | 0.623 |
| ReNet | 0.171 | 0.765 | 0.096 |
| BeFree | 0.105 | 0.688 | 0.057 |

**No Ambiguity Consensus**

| | F-score | Precision | Recall |
|---|---|---|---|
| DTMiner | 0.8 | 0.753 | 0.852 |
| GeDex | 0.668 | 0.78 | 0.584 |
| ReNet | 0.129 | 0.833 | 0.07 |
| BeFree | 0.096 | 0.717 | 0.051 |

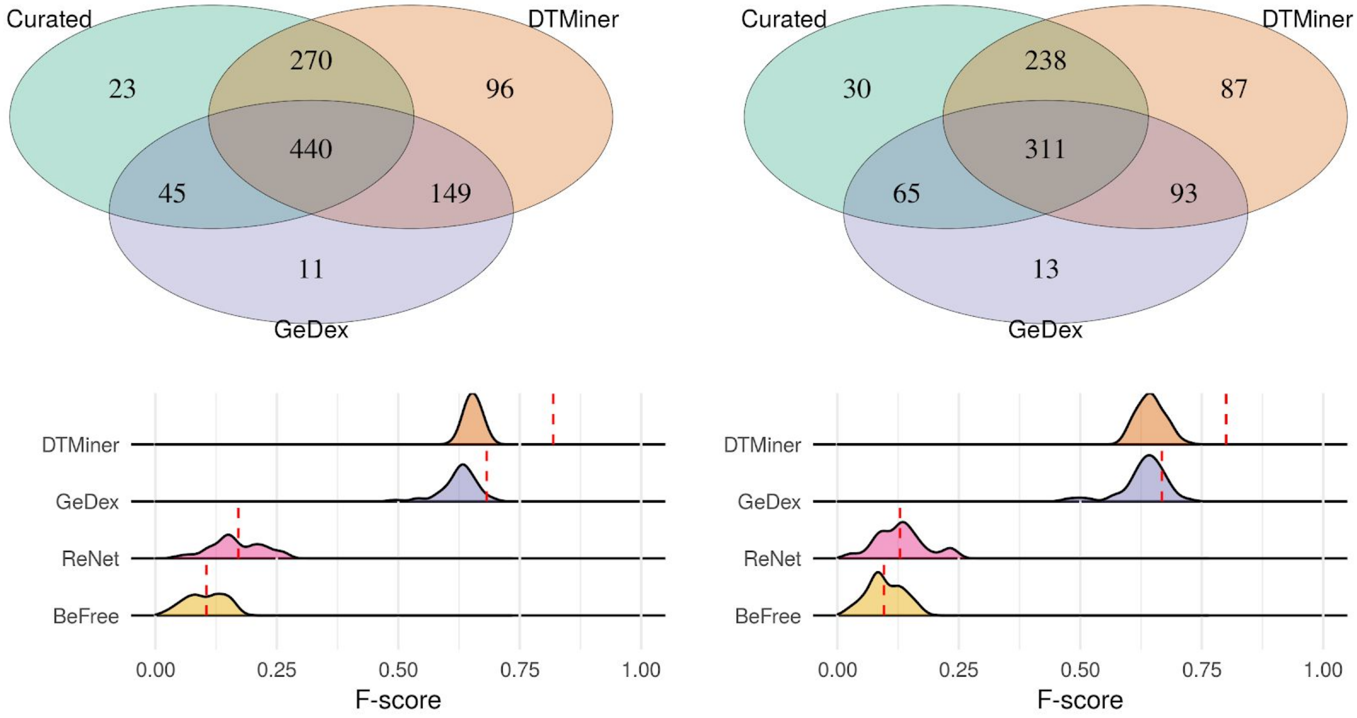


**Figure S5.** GeDex validation and benchmarking. Here we show complementary results of Figure 3, that were obtained with the Majority Consensus (left column) and the No Ambiguity Consensus (right column)

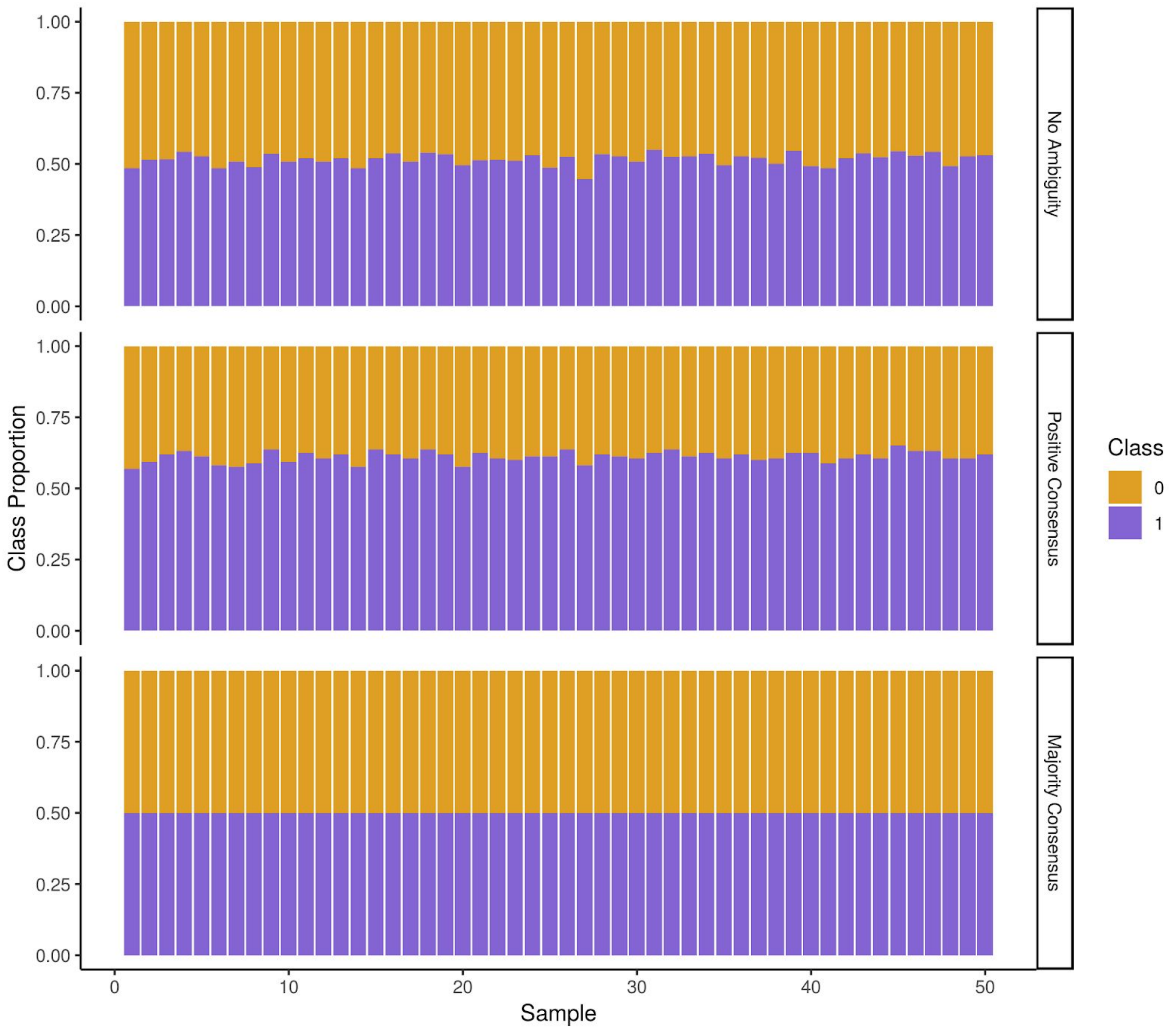


**Figure S6.** Class distribution in manually-curated corpus samples. Each sample contains the same interactions in each Consensus type. The class distribution in the complete manually-curated corpus can be seen in the column marked as **CR** (complete reference), which shows a high imbalance class.
